# Proteome signatures of reductive stress cardiomyopathy

**DOI:** 10.1101/2021.09.13.460105

**Authors:** Sini Sunny, Cynthia L. David, Krishna Parsawar, Dean P. Jones, Namakkal S. Rajasekaran

## Abstract

Nuclear factor erythroid 2-related factor 2 (NRF2), a redox sensor, is vital for cellular redox homeostasis. We reported that transgenic mice expressing constitutively active Nrf2 (CaNrf2-TG) exhibit reductive stress (RS). In this study, we identified novel protein biomarkers for RS-induced cardiomyopathy using Tandem Mass Tag (TMT) proteomic analysis in heart tissues of TG (CaNrf2-TG) and non-transgenic (NTg) mice at 6-7 months of age (N= 4/group). A total of 1105 proteins were extracted from 22544 spectra. Of note, about 560 proteins were differentially expressed in TG vs. NTg hearts, indicating a global impact of RS on myocardial proteome. From a closer analysis of the proteome datasets, we identified over 32 proteins that were significantly altered in response to RS. Among these, 20 were upregulated and 12 were downregulated in the hearts of TG vs. NTg mice, suggesting that these proteins could be putative signatures of RS. Scaffold analysis revealed a clear distinction between TG vs NTg hearts. Of note, we observed several proteins with redox (#185; cysteine residues), NEM-adducts (#81), methionine-loss (#21) and acetylation (#1) modifications in TG vs. NTg hearts due to chronic RS. The majority of the differentially expressed proteins (DEPs) that are significantly altered in RS mice were found to be involved in stress related pathways such as antioxidants, NADPH, protein quality control (PQC), etc. Interestingly, proteins that were involved in mitochondrial respiration, lipophagy and cardiac rhythm were dramatically decreased in TG hearts. Of note, we identified the glutathione family of proteins as the significantly changed subset of the proteome in TG heart. Surprisingly, our comparative analysis of NGS based transcriptome and TMT-proteome indicated ∼50% of the altered proteins in TG myocardium was found to be negatively correlated with their transcript levels. Modifications at cysteine/NEM-adducts (redox), methionine or lysine residues in multiple proteins in response to chronic RS might be associated with impaired PQC mechanisms, thus causing pathological cardiac remodeling.

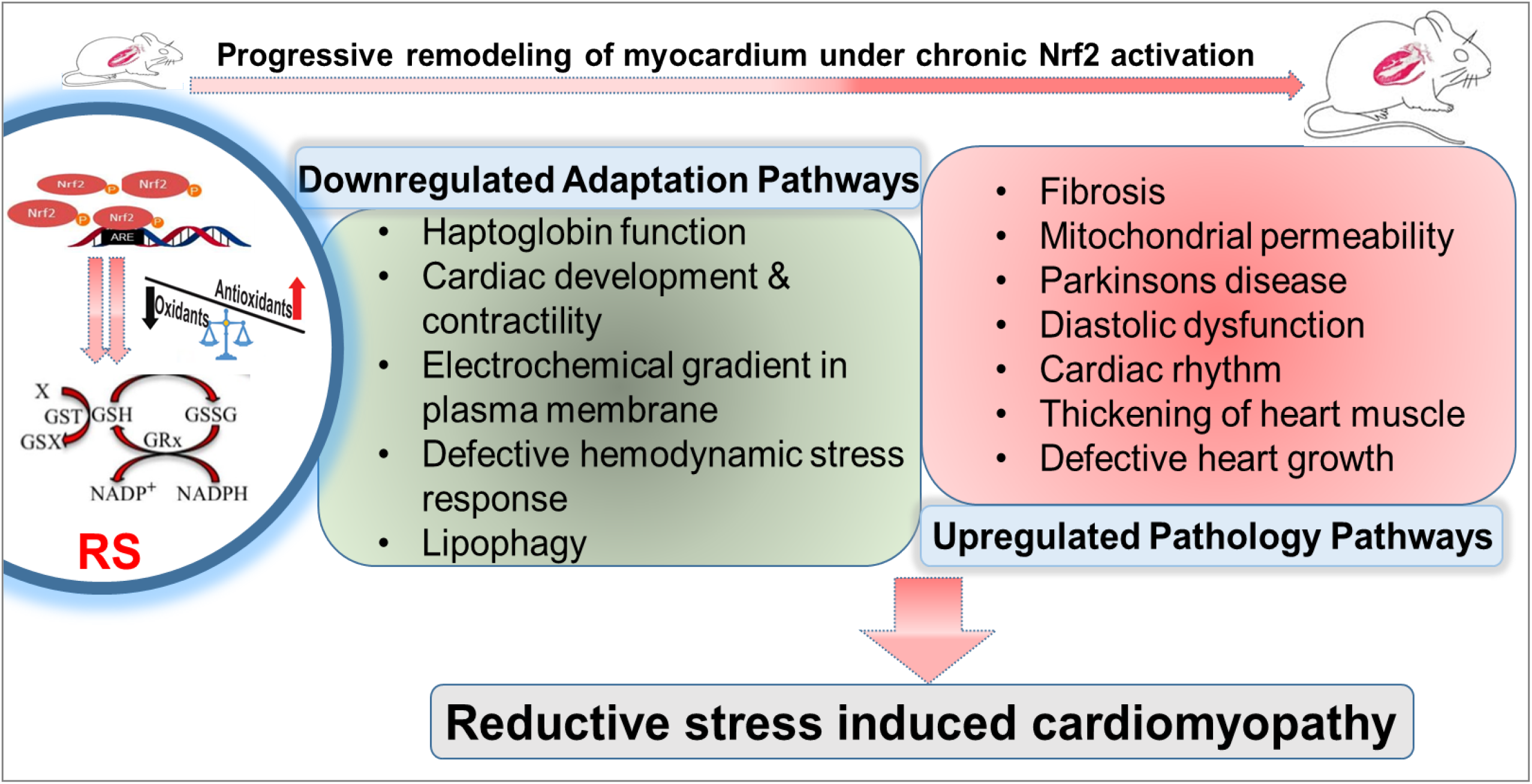
Graphical Abstract.

## 1. Introduction

A real-time action of Nrf2 in response to stress is vital to preserve the redox homeostasis in cells and tissues, but its activation under unstressed settings could tilt the redox balance towards the reductive arm, leading to reductive stress (RS). Cardiac-specific transgenic mouse models expressing constitutively active Nrf2 (caNrf2-TG) develop RS and cause pathological cardiac remodeling (1). We recently reported that caNrf2-TG mice exhibit eccentric cardiac hypertrophy with increased ejection fraction and progressive diastolic dysfunction (1). Many studies from other investigators and we revealed global changes in the transcriptome and pathophysiological processes in heart, skeletal muscle, metabolism, cancer, and neurodegeneration under reductive and hyper-reductive conditions (2-8). Likewise, caNrf2-TG mice displayed a unique transcriptome profile that is believed to drive the pathological remodeling of the myocardium in a chronic setting (9, 10). Nonetheless, the proteome alterations associated with RS and subsequent cardiac remodeling are unknown.

From the past decade, many other studies indicated that a reductive intracellular environment (i.e., RS) might be challenging to normal physiological signaling processes due to lack of basal reactive oxygen and nitrogen species (ROS/RNS) (11-14). Particularly, ROS/RNS are necessary for cell proliferation and differentiation during embryonic development, regeneration of stem cells/tissues and healing of damaged tissues, etc. (15-19). We recently reported that either acute or chronic reductive stress conditions impairs the regeneration of myoblasts and neuroblastoma cells (2, 3). Notably, RS-induced neurodegeneration is driven by misfolding and aggregation of proteins (3), but the mechanisms are unknown.

In the present study, we aimed to understand the impact of RS on global proteome changes in the myocardium using tandem mass tagging (TMT)-based proteomics. We performed TMT-proteomic analysis of the myocardial cytosolic proteins isolated from non-transgenic (NTG) and transgenic (caNrf2-TG) RS mice. We hypothesized that Nrf2 mediated RS will alter the myocardial redox proteome and perturb the physiological processes.

## 2. Materials and Methods

### 2.1. Animals

Heart-specific constitutively active Nrf2 (CaNrf2) transgenic (TG) and non-transgenic (NTG) mice (n=4/group) at 6 months of age were used for analyzing the myocardial proteome. Animals were maintained under hygienic condition with a 12 h day/night cycle and fed with standard animal chow and water *ad libitum*. All studies were conducted in accordance with the Guide for Care and Use of Laboratory Animals developed by the National Research Council at the National Institutes of Health (NIH). The Institutional Animal Care and Use Committee (IACUC#14-10160) at the University of Alabama at Birmingham has approved the study.

### 2.2. Sample preparation for Tandem Mass Tag (TMT) proteomic analysis

TMT-6plex reagents from Thermo were used to isotopically label peptides following reduction and alkylation of 60ug mouse heart lysate by following the vendor’s protocols. Heart tissues were harvested from non-transgenic (NTG) & caNrf2-TG-High (TGH) (N=4 mice/group) after perfusing with N-ethylmaleimide (NEM; 10 μM). Heart tissue homogenates were prepared in RIPA buffer, and the cytosolic proteins were separated by centrifugation (5000 rpm, for 7 minutes at 4°C). Protein concentration was determined with a BCA kit (Bio-Rad, USA) according to the manufacturer’s instructions. After trypsin digestion, peptide was desalted by Strata-X C18 SPE column (Phenomenex, Torrance, CA, USA) and vacuum-dried. Peptide was reconstituted in 0.5 M TEAB and processed according to the manufacturer’s protocol for a TMT (Tandem Mass Tag) kit (ThermoFisher Scientific, Waltham, MA, USA). Briefly, one unit of TMT reagent was thawed and reconstituted in acetonitrile. The peptide mixtures were then incubated for 2 h at room temperature and pooled, desalted, and dried by vacuum centrifugation (20).

### 2.3. LC MS/MS Method and Data Searching

LC-MS/MS analysis was performed on a Q Exactive Plus mass spectrometer (Thermo Fisher Scientific, San Jose, CA) equipped with an EASY-Spray nanoESI source. Peptides (500ng) were eluted from an Acclaim Pepmap 100 trap column (75 micron ID x 2 cm, Thermo Scientific) onto an Acclaim PepMap RSLC analytical column (75 micron ID × 25 cm, Thermo Scientific using a 3-25% gradient of solvent B (acetonitrile, 0.1% formic acid) over 150 min, 25-50% solvent B over 20 min, 50-70% of solvent B over 10 min, 70-95% B over 10 min then a hold of solvent 95% B for 20 min, and finally a return to 3% solvent B for 10 min. Solvent A consisted of water and 0.1% formic acid. Flow rates were 300 nL/min using a Dionex Ultimate 3000 RSLCnano System (Thermo Scientific). Data dependent scanning was performed by the Xcalibur v 4.0.27.19 software (21) using a survey scan at 70,000 resolution scanning mass/charge (m/z) 375-1400 at an automatic gain control (AGC) target of 3e6 and a maximum injection time (IT) of 50 msec, followed by higher-energy collisional dissociation (HCD) tandem mass spectrometry (MS/MS) at 32nce (normalized collision energy), of the 11 most intense ions at a resolution of 35,000, an isolation width of 1.2 m/z, an AGC of 1e5 and a maximum IT of 100 msec. Dynamic exclusion was set to place any selected m/z on an exclusion list for 30 seconds after a single MS/MS. Ions of charge state +1, 7, 8, >8, unassigned, and isotopes were excluded from MS/MS (22).

### 2.4. Protein identification

MS and MS/MS data were searched against the amino acid sequence of the Uniprot mouse protein database and a common contaminant protein database (e.g., trypsin, keratins; obtained at ftp://ftp.thegpm.org/fasta/cRAP) using Thermo Proteome Discoverer v 2.4.0.305 (Thermo Fisher Scientific). MS/MS spectra matches considered fully tryptic peptides with up to 2 missed cleavage sites. Variable modifications considered were methionine oxidation (15.995 Da), and cysteine carbamidomethylation (57.021 Da). Proteins were identified at 95% confidence with XCorr score cut-offs (23) as determined by a reversed database search.

### 2.5. Quantitative data analysis

The protein and peptide identification results were further analyzed with Scaffold Q+S v 4.11.1 (Proteome Software Inc., Portland OR), a program that relies on various search engine results (i.e.: Sequest, X!Tandem, MASCOT) and which uses Bayesian statistics to reliably identify more spectra (22). Protein identifications were accepted that passed a minimum of two peptides identified at 95% protein and peptide confidence levels. Quantitative values were generating using the Proteome Scaffold Q module. Acquired intensities in the experiment were globally normalized across all acquisition runs. Individual quantitative samples were normalized within each acquisition run. Intensities for each peptide identification were normalized within the assigned protein. The reference channels were normalized to produce a 1:1 fold change.

The data, analytic methods, and study materials will be made available to other researchers for purposes of reproducing the results or replicating the procedure. The mass spectrometry proteomics data have been deposited to the ProteomeXchange Consortium (http://proteomecentral.proteomexchange.org) via the PRIDE partner repository.

### 2.6. Next generation RNA sequencing

Total RNA was isolated from TG and NTG mice (n=4/group) at 6 months of age using the RNeasy Mini Kit (Qiagen, Cat.74106) according to the manufacturer’s instructions. The purity of RNA was confirmed using Bio-analyzer and intact poly(A) transcripts were purified from total RNA using oligo(dT) magnetic beads and mRNA sequencing libraries were prepared with the TruSeq Stranded mRNA Library Preparation Kit (Illumina, RS-122-2101, RS122-2102) (9).

### 2.7. Immunoblotting

Independent validation of some of the MS data was carried out using protein specific antibodies. Heart tissues from NTG and caNrf2 TG mice (n = 4/group) at 6 months of age were homogenized in a cytosolic extraction buffer (10 mM HEPES, 10 mM KCl, 0.1 mM EDTA, 0.5 mM MgCl2, with freshly prepared 0.1 mM phenyl methylsulfonyl fluoride (PMSF), 1 mM dithiothreitol, and 1% Triton X-100, pH 7.9) and centrifuged at 5000 rpm for 5–6 min. Protein concentrations were determined with the Bradford reagent, and equal amounts of protein were resolved on 10%–12% sodium dodecyl sulfate– polyacrylamide gel electrophoresis. Proteins were then transferred to polyvinylidene difluoride membranes (EMD Millipore Corp., Billerica, MA) and blocked-in tris-buffered saline-Tween 20 (TBST) containing 5%–10% nonfat dry milk or BSA. Membranes were then incubated overnight at 4°C with their respective primary antibodies (1%–2% BSA in TBST) followed by horseradish peroxidase IgG (Vector Laboratories)-conjugated secondary antibody (anti-rabbit and anti-mouse, 0.2% BSA in TBST) incubation followed by chemiluminescence-based detection (Pierce, Rockford, IL) with an Amersham Imager 600 (GE Healthcare Life Sciences, Chicago, IL) (24).

### 2.8. Bioinformatics tools

Differentially expressed proteins (DEPs) were then plotted for heat maps based on fold change with R studio package “pheatmap” (Version 1.2.5033). PCA plot was plotted using ClustVis (https://biit.cs.ut.ee/clustvis/). We performed Protein Analysis THrough Evolutionary Relationships (PANTHER) (pantherdb.org) to predict the functions of uncharacterized proteins in TMT proteome, based on their evolutionary relationships. Also, we carried out STRING analysis (https://string-db.org/), to predict protein-protein interactions - direct (physical) and indirect (functional) associations between the proteins analyzed in the scaffold software.

### 2.9. Statistics

*Proteomic data*: P value (<0.05) for differentially expressed proteins were calculated using Bonferroni Corrected Mann–Whitney tests (Scaffold 4.0). *Western blotting*: Mean ± standard error is shown (n = 4 hearts per experimental group). Comparisons were performed by one-way ANOVA. P < 0.05 was considered statistically significant.

## 3. Results

### 3.1. Identification of proteins in RS (TG) hearts

Quantitative proteomic analysis was performed on NTG and TG heart tissues using six plex-tandem mass tag (TMT) labeling. A total of 22544 quantitated spectra (out of 24372 threshold spectra) with 95% minimal False Discovery Rate (FDR) representing 1105 proteins (889 clusters) were identified in the TG heart tissues (**Fig.1A**). Comparative spectral profiling based on log fold change identified 560 differentially expressed proteins (DEPs) with a significant log2 fold change (FC) ≥1.2, 168 DEPs at FC≥1.5, 32 DEPs at FC≥2, 3 DEPs at FC ≥4 and 1 DEP at FC≥8 in TG *vs* NTG mouse hearts (**Fig.1B**). Principle compound analysis using 2D scatter plot (PCA plot) revealed distinct clustering of TG (caNrf2-TG) and NTG samples (**Fig.1C**), suggesting the clear role impact of RS on myocardial proteome. At 95% protein threshold confidence level, significant difference between proteome distributions in each group was observed. R studio analysis distributed the whole DEPs into distinct bins based on log2 fold change (**Fig.1D)**.

**Figure 1:**
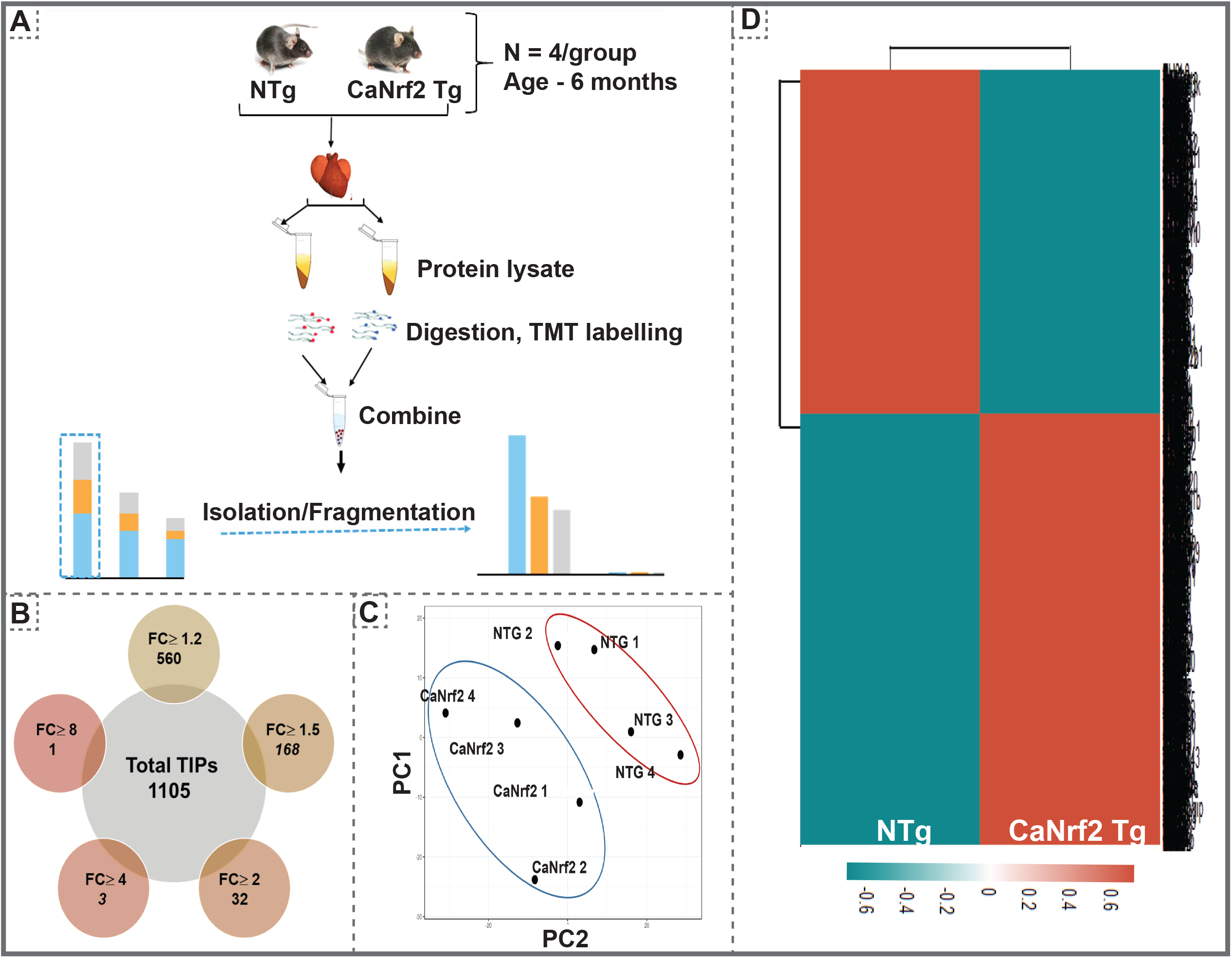
(A) Overall methodology adopted for Tandem Mass Tagged LC-MS/MS analysis. Heart tissues were harvested from non-transgenic (NTG) & caNrf2-Tg (N=4 mice/group) and protein concentration was determined in homogenates with BCA kit (Bio-Rad, USA). After trypsin digestion, peptide was reconstituted in 0.5 M TEAB and processed for TMT tagging (Tandem Mass Tag kit, ThermoFisher Scientific). LC-MS/MS analysis was performed on a Q Exactive Plus mass spectrometer (Thermo Fisher Scientific) equipped with an EASY-Spray nanoESI source. MS and MS/MS data were searched against the amino acid sequence of the Uniprot mouse protein database using Thermo Proteome Discoverer v 2.4.0.305 (Thermo Fisher Scientific). The protein and peptide identification results were further analyzed with Scaffold Q+S v 4.11.1 (Proteome Software Inc.). (B) Venn diagram showing the number of Differentially Expressed Proteins (DEPs) based on different fold change (CaNrf2 Tg vs NTg) identified in TMT proteome software (C) Prinicipal Component Analysis (PCA) Plot generated using Total Identified Proteins (TIPs) showed segregation of NTG and CaNrf2 Tg as distinct groups. (D) Global heat map generated using R studio for TIPs (1105) proteins identified in TMT analysis.

### 3.2. Gene ontology and pathway analysis

To gain further insight into the enriched pathways associated with the RS proteome in the heart, core proteome was submitted for panther (**Fig.2A)** and string analysis (**Fig.2B)**. Functional classification using Panther showed an enrichment of metabolite interconversion enzyme family (i.e., oxidoreductase family), which are directly or indirectly contributing to myocardial health, in TG hearts. Among the metabolic enzymes, oxidoreductases were 80% enriched in TG hearts (**Fig.2A)**. Mainly, the protein targets of Nrf2, NQO1, GPX1, GPX4, SOD2 were detected by Panther in CaNrf2-TG hearts validates the signature for RS. Other proteins that are directly or indirectly associated with RS proteome are listed in **Fig.2A**. Furthermore, String analysis clustered the proteome core into different groups, based on the physical protein association, with NADPH as the most enriched network. Based on functional/physical protein associations and kmeans clustering method, string segregated the proteins into three clusters, which are tightly enriched and regulated by ubiquitin, GSR and stress protein families (**Fig.2B)**.

**Figure 2.**
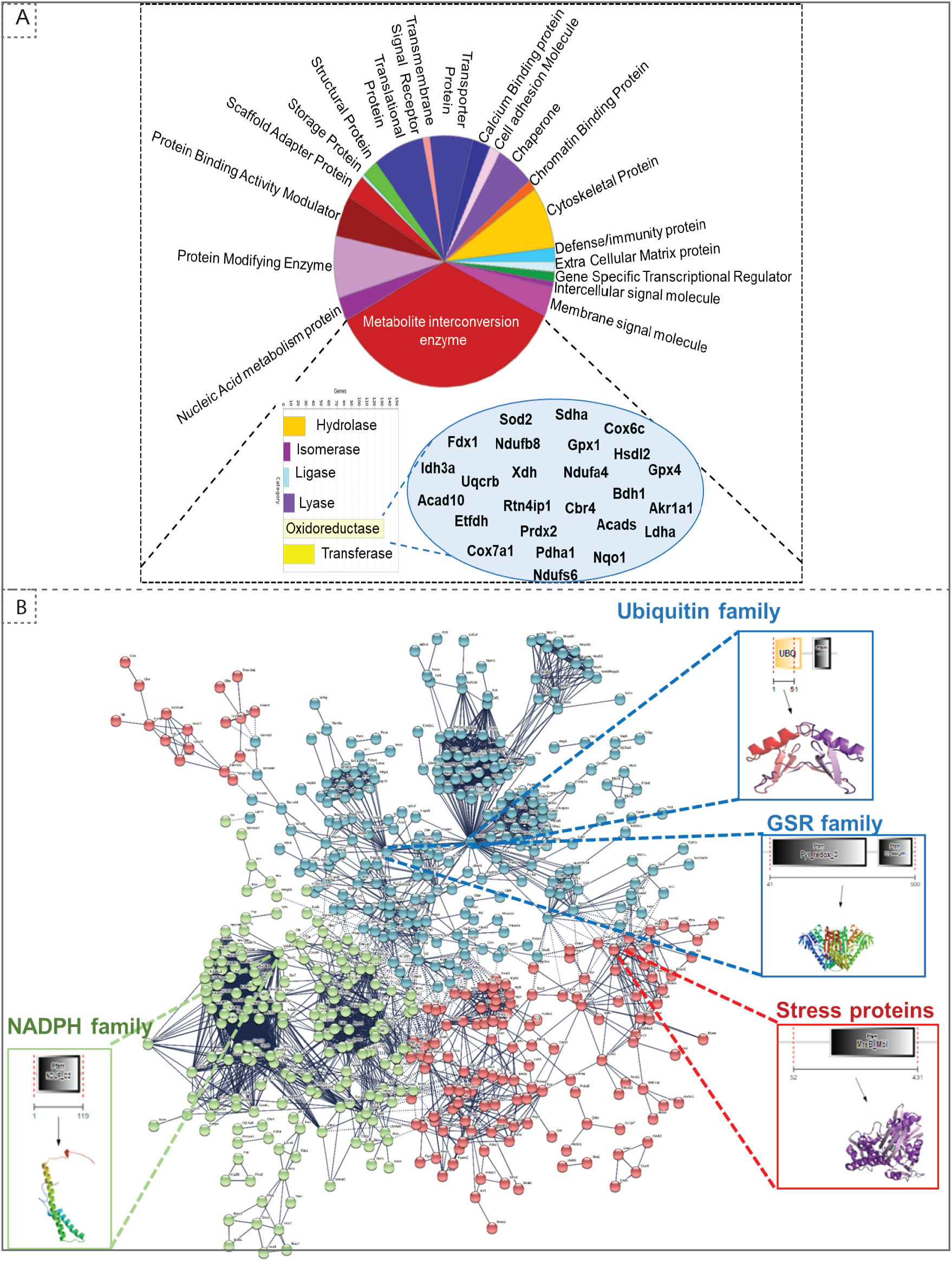
Gene ontology analysis for DEPs identified in TMT analysis. (A) Protein Analysis THrough Evolutionary Relationships (PANTHER) analysis (http://pantherdb.org/) for biological function distributed the TIPs into different metabolic categories. 25% of the proteome function grouped under metabolite interconversion category, with oxidoreductase as the top upregulated enzyme functional category. Those proteins identified under oxidoreductase group is shown in oval shape. (B) STRING analysis (https://string-db.org/) for predicted protein-protein interactions showed different subset of gene interaction network with ubiquitin, GSR, stress proteins, NADPH as the top enriched pathway in TG heart. The network has been refined using K means clustering method with a highest confidence cut off 0.900. Protein’s nodes are colored automatically based on the number of clusters opted.

### 3.3. Peptide modifications in RS hearts

Using unsupervised clustering, we identified 32 DEPs (FC≥2) including GSTA1 and GSTA3 showing a highest enrichment in TG hearts. Other proteins like PRDX6, TALDO1, BLVRD, BIN1, GSTM1 and PIR were recognized as the second big enrichment (**Fig.3A)**. Of note, we identified several post-translational modifications such as oxidation (185), N-ethylmaleimide (NEM; 120), methionine loss (21), acetylation (1) and methionine loss + acetylation (1) in comparison with NTG (**Fig.3B**). It is interesting that these observations are associated with the chronic reductive environment. Future studies warrant investigations on the conformational changes in each protein and the molecular mechanisms coupled with these alterations.

**Figure 3.**
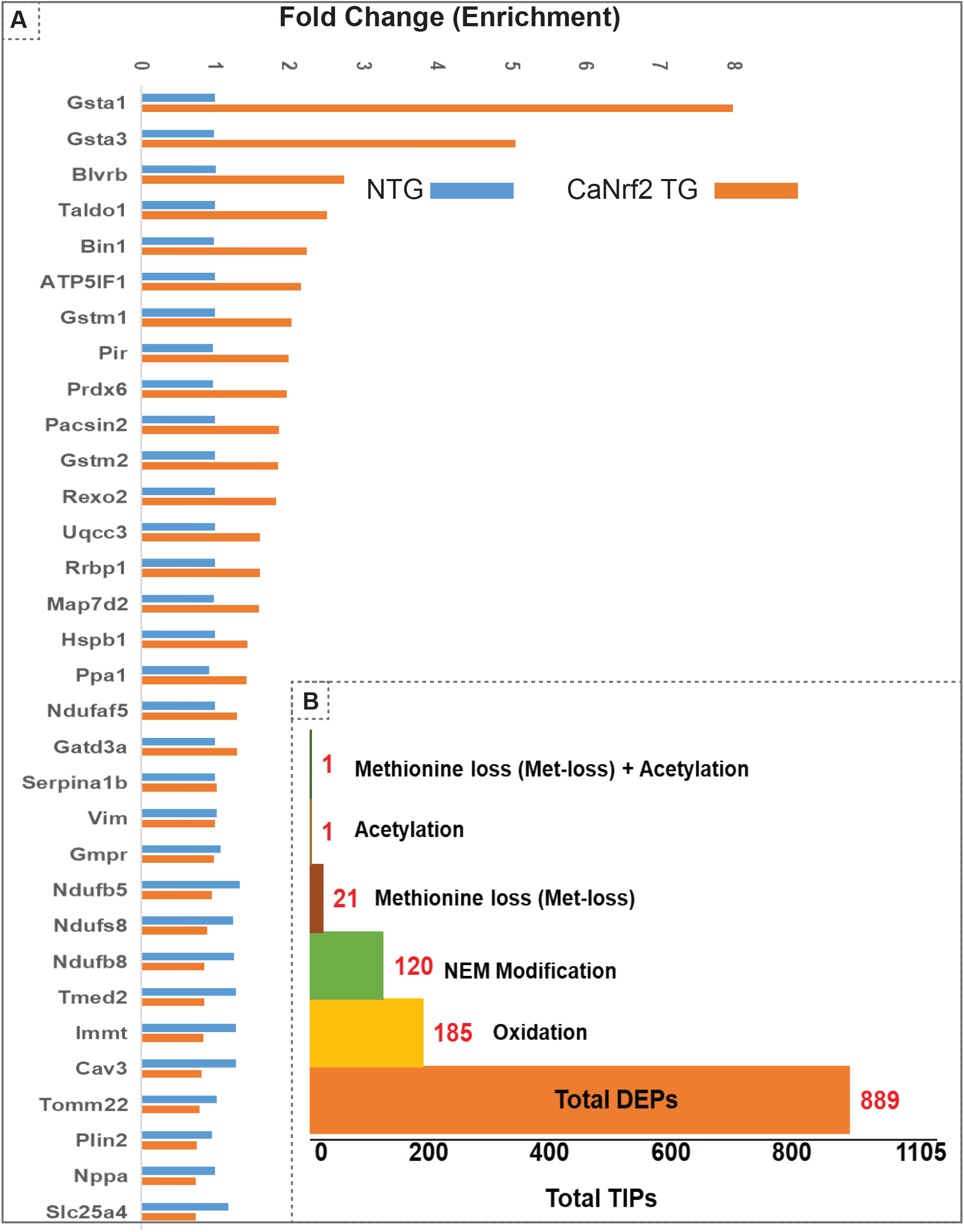
(A) Unsupervised clustering of total identified proteins showed 32 DEPs at 2 fold change with Gsta1, Gsta2, Blvrd, Taldo1, Bin1, Atp5lf1, gstm1,Pir and Prdx6 as the top upregulated proteins in CaNrf2 hearts (B) Spectrum view by proteome scaffold software identified different types of modification in differentially expressed proteins (889) with 185 proteins having oxidation,120 with NEM modification, 21 with loss of methionine peptide, 1 with acetylation, 1 with acetylation and 1 peptide with both loss of methylation and acetylation.

### 3.4. Putative indicators of reductive stress

Interestingly, among the DEPs (up- or down-regulated), we noticed some of them were intact while others had modifications at specific amino acid residues in the RS hearts **(Fig.4A**). A list of top identified proteins with their modifications in peptides and the amino acid residues is shown **Fig.4B**. Several proteins had modifications in cysteine, methionine, and lysine residues. Glutathione metabolism is tightly regulated and has been implicated in myocardial redox signaling (25). Interestingly, we observed glutathione enzyme family and other related proteins were robustly upregulated in TG (FC>2.7) in comparison to NTG hearts (**Fig.4C**). Spectrum analysis of GSTA1, GSTA2, GSTA3, GSTA4, GSTM1, GSTM3 and GSTM4 recognized cysteine, lysine, and methionine modifications in their peptide fragments (Fig.SX). In fact, we validated and confirmed the expression of some of these proteins using immunoblotting (**Fig.4D**).

**Figure 4.**
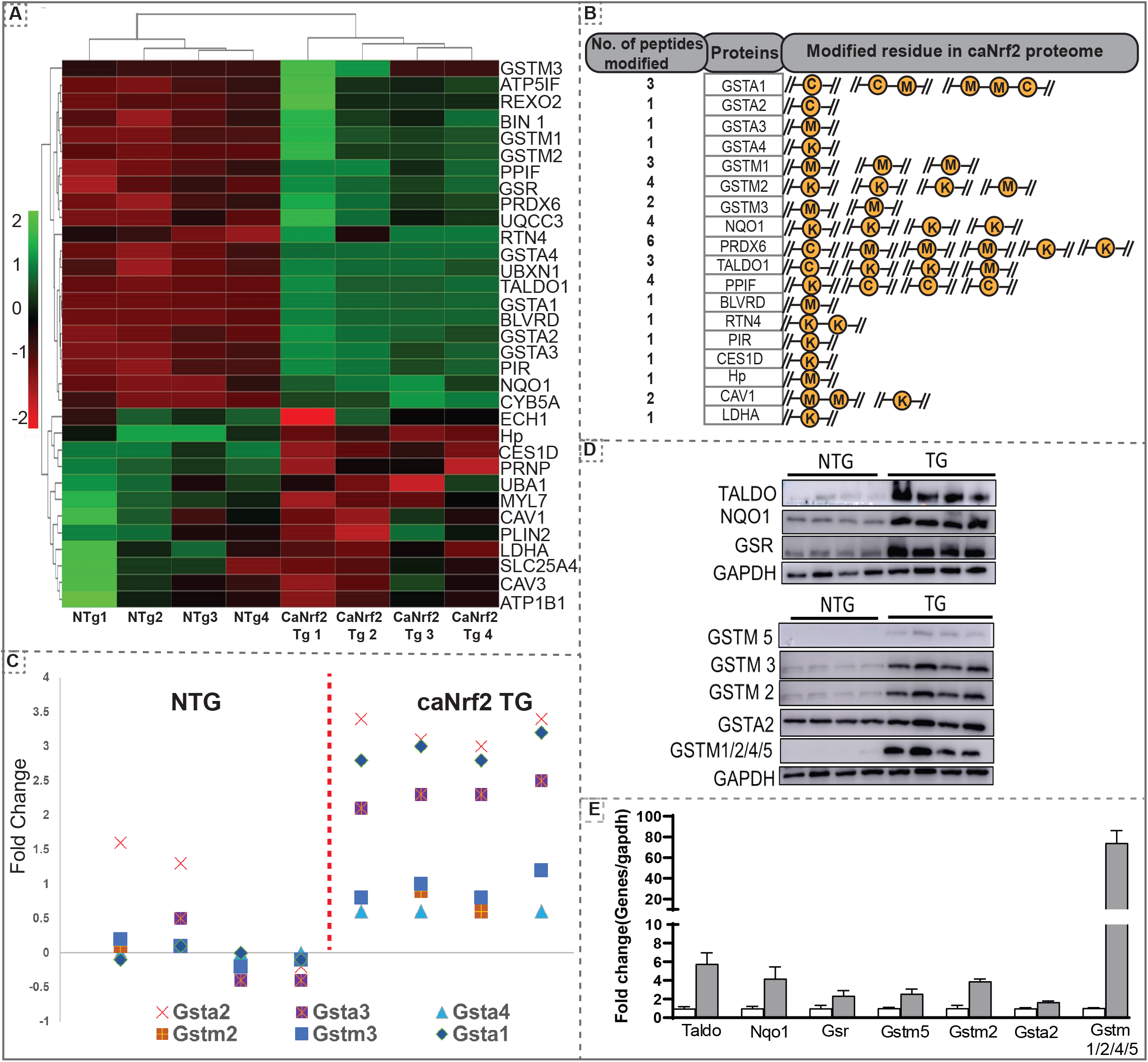
(A) CaNrf2 proteome heat map using R studio for highly upregulated proteins (based on log2 FC) identified in scaffold software (B) Highly upregulated proteins are associated with amino acid modifications in peptide(s). Data showing the number of peptides modified in each protein and modified residue in each peptide. A representative peptide is shown in yellow color. (C) TMT proteome identified GSR family of proteins as top enriched in CaNrf2 hearts in comparison with NTG. Significantly changed proteins are Gsta2, Gsta3, Gsta4, Gstm2, Gstm3 and Gsta1 (based on log2 fold change ratio). (D) Immunoblot validation for the selected proteins in Tg vs NTG hearts. (E) Densitometry analysis using ImageJ is shown.

### 3.5. Comparative analysis of transcription (NGS-RNA-Seq) and translation (TMT-Proteomics) of protein targets in the RS myocardium

Comparison of RNA sequencing data [Next Generation Sequencing (NGS)] with proteomic data (TMT) from CaNrf2-TG hearts showed surprising results. About 50% of the proteins do not match with the quantitative changes of their genes (mRNA; **Fig.5**). Out of 104 commonly found gene products in NGS (RNA) and respective translational products (proteins by TMT) at a FC>1.5, the levels of 50 proteins are not consistent with their RNA levels (either up or down regulated) (**Fig.6A**). The proteins that are steadily expressed according to their respective mRNA levels are classified as “transcription-sync proteins” (**Fig.6B**) and the ones that do not match with their mRNA levels are termed as “transcription-nonsync proteins” (**Fig.6C**). Among the transcription-sync proteins, 16 (MAP7D2, BLVRD, BIN1, MYL4, DAP, TALDO1, PIR, NPPA, GGT5, PRDX6, NDUFAF5, GSTM2, RTN4, GSR, ALD, and BRACL) with marginally higher RNA levels were statistically insignificant (r=0.194). As expected, changes in Nrf2 targeted antioxidant proteins are statistically significant (NQO1, GSR, GSTA1, GSTA3, GSTA4, and GSTM1) and grouped under “transcription-sync proteins”. Further, 9 of the “transcription-nonsync proteins” (ACADVL, ATP5PB, CES1D, CMBL, COX7A1, CPT1B, FAH, GSTK1, and HADHA) showed high protein levels albeit their significantly down regulated mRNA expression (**Fig.6C**). A strong positive correlation (r=0.955) was observed among 38 of transcription-sync proteins (**Fig.6D & E**), but a weak positive correlation (r=0.194) for 16 transcription-sync proteins (**Fig.6F**). Transcription-nonsync proteins showed a negative correlation (**Fig.6G**).

**Figure 5.**
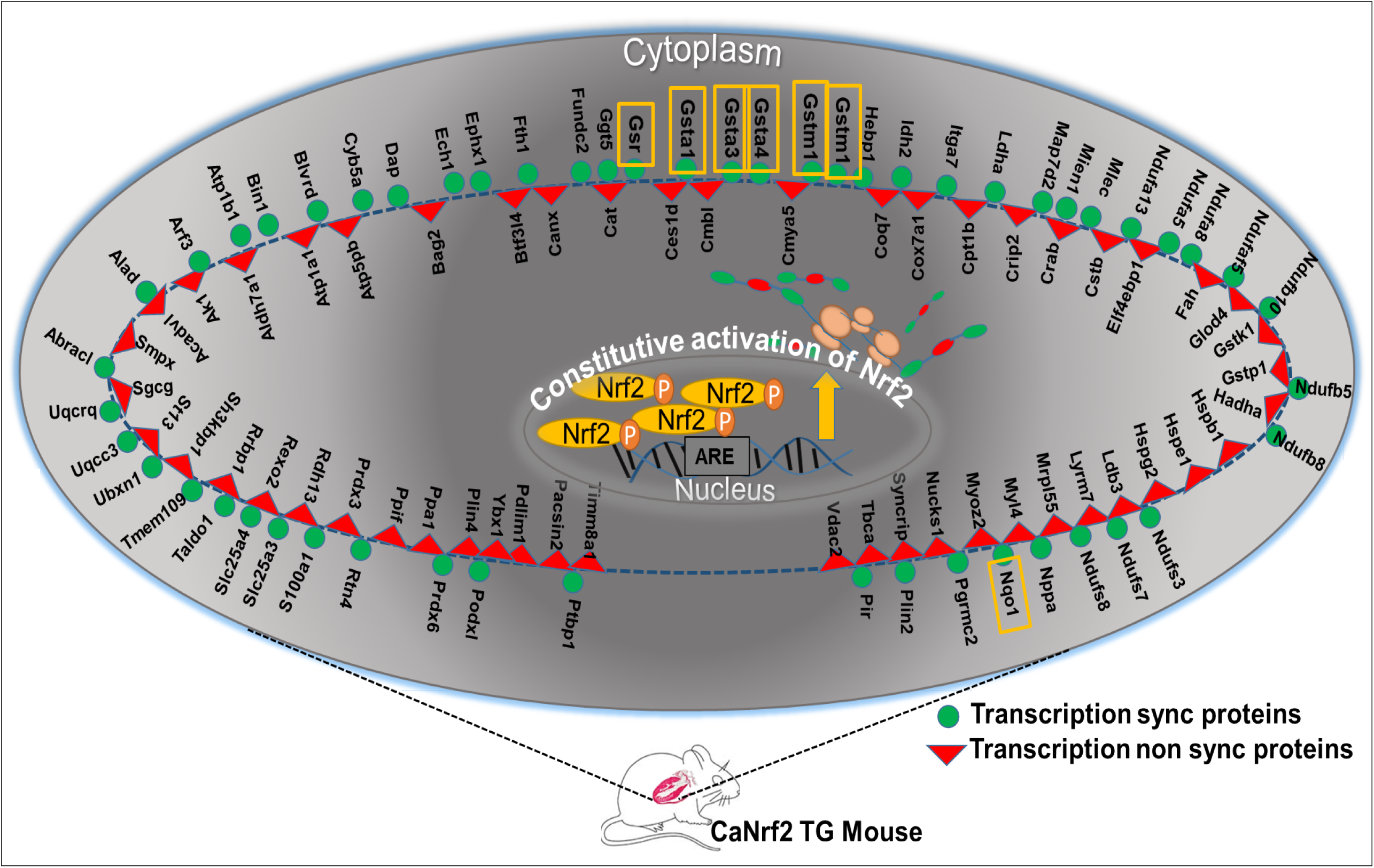
Comparative analysis between RNA seq and TMT proteome data in CaNrf2 hearts. About 50% of the proteins do not match with the quantitative changes of their genes. The proteins that are steadily expressed according to their respective mRNA levels are classified as “transcription-sync proteins” (Fig.6B) and the ones that do not match with their mRNA levels are termed as “transcription-nonsync proteins”. Nrf2 targeted antioxidant proteins are statistically significant (NQO1, GSR, GSTA1, GSTA3, GSTA4, and GSTM1) and grouped under “transcription-sync proteins” and highlighted in yellow boxes.

**Figure 6.**
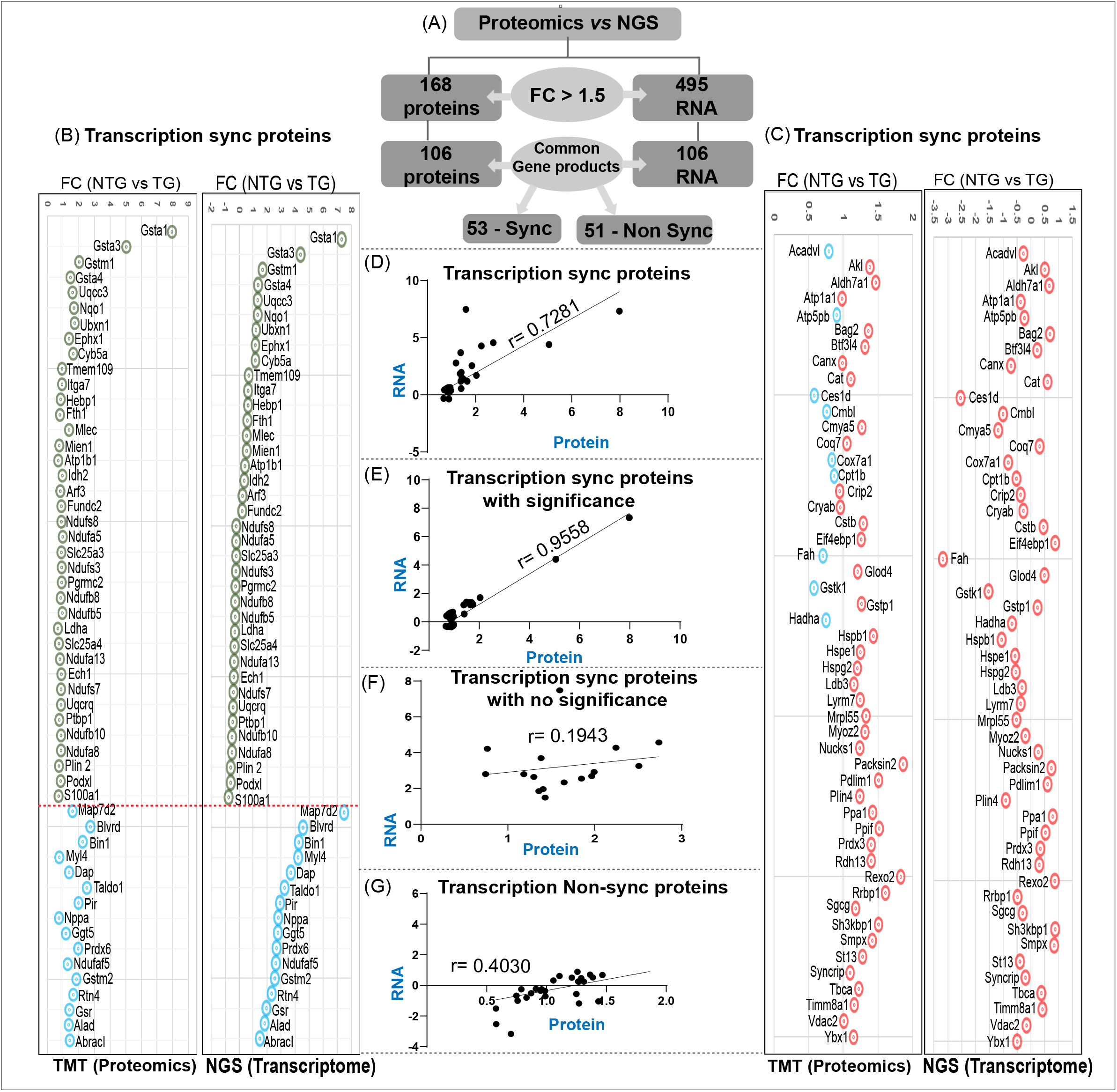
Correlation between transcription-sync proteins and transcription-non sync proteins in caNrf2 Tg heart. (A) Out of 104 commonly found gene products in NGS (RNA) and respective translational products (proteins by TMT) at a FC>1.5, the levels of 50 proteins are not consistent with their RNA levels (either up or down regulated). Proteins that are steadily expressed according to their respective mRNA levels are classified as “transcription-sync proteins” (B) and the ones that do not match with their mRNA levels are termed as “transcription-nonsync proteins” (C). A strong positive correlation (r=0.955) was observed among 38 of transcription-sync proteins (D & E), but a weak positive correlation (r=0.194) for 16 transcription-sync proteins (F). Transcription-nonsync proteins showed a negative correlation (G).

## 4. Discussion

Constitutive activation of Nrf2 under normal (unstressed) condition is a key propagator for RS. Our previous data on CaNrf2 transgenic mice (with RS) revealed a distinct reductive transcriptome in these hearts (1). To elucidate the proteomic profiling of RS hearts, we employed tandem mass tagging (TMT) based mass spec analysis and distinguished the differentially expressed proteins (DEPs). Proteomic profiling and computational pathway analyses provide both over-all and specific information associated with RS pathology. TMT LC-MS/MS analysis identified a total of 1105 proteins across the two groups-caNrf2-TG and NTG, which were clearly segregated in a PCA plot. Distinct bins clustered in heat map analysis for TG and NTG also validates the reductive stress dependent protein profile in TG mouse myocardium.

To gain evidence for reductome-signature in TG hearts, gene ontology pathway analysis was carried out by panther and string analysis (26, 27). Panther analysis classify the proteins on the basis of biological function and pathways (28). Panther categorized oxidoreductase enzymes as top enriched pathway along with other pathways related to structural/functional adaptions during the cardiac development. String analysis was used to predict direct (physical) and indirect (functional) association between the proteins identified through scaffold (29). Clustering and enrichment of NADPH associated proteins, suggesting a possible development of reductive environment in the TG hearts. To refine the closely interacting protein families, Kmeans-based clustering was performed. The functional analysis showed three different clusters with enrichment of ubiquitin, GSR, NADPH family and stress proteins in the core proteome. Unbiased refining of the whole proteome with Panther and String showed high enrichment of metabolic enzymes crucial for redox homeostasis and proteins involved in redox signaling pathways. These results support the distinct reductome (reductive stress proteome) signatures in caNrf2-TG hearts. Among the DEPs responding to RS, several proteins were modified in the TG when compared to NTG hearts. Modifications like N-ethylmaleimide (NEM), oxidation, methylation and acetylation were obvious in TG/RS hearts. Interestingly, most of the 185 oxidized peptide fragments were exclusively detected in mass spectrum sequences of TG hearts (**Fig.3B**).

We also observed several upregulated proteins with no modifications and down regulated proteins with/without modifications in CaNrf2 cardiac proteome. Noteworthy, we identified glutathione and its related family proteins were robustly enriched in TG hearts. The capacity to recycle GSH makes the glutathione system pivotal to the antioxidant defense mechanism and preserves the cellular thiols (30). Proteome analysis revealed multiple modifications in more than one peptide of GSH family proteins (oxidation of cysteine in GSTA1 and GSTA2). While GSTA3 displays methionine oxidation, GSTA4 has lysine oxidation in one of the peptide fragments. However, in GSTM1, GSTM2 and GSTM3, both methionine and lysine residues were modified. Modifications in the cysteine residues of these GST family enzymes might impair their kinetics in the RS myocardium. Noxious effects of such modifications in free cysteine residues that are in queue for t-RNA selection may dramatically alter the translation process under RS.

More interestingly, comparison of RNA-seq data (NGS) with TMT proteomic data in CaNrf2 hearts elucidated that, a RS myocardium is not following the quantitative omics. We compared and classified the gene products (RNA and proteins) based on their expression levels as transcription-sync and transcription-nonsync proteins. Interestingly, we found that transcription versus translation process is not fully synchronized in the RS myocardium. Observed changes in the expression of proteins in response to RS might be primarily caused by (a) direct trans-activation of target genes of Nrf2, (b) chronic impact of RS, (c) RS-mediated posttranslational changes and (d) positive- or negative-feedback of protein synthesis rate on the transcription of the respective gene(s). We also hypothesize that changes in the translational efficiency are caused, in part, by aggregation of proteins which might trigger a feedback inhibition of the translation, or due to the modified amino acid residues, which alter the ribosomal attachment and decoding of mRNA, which results in synthesis of partial or over abundant transcript during the translation (31-34). Moreover, modified peptides observed in the caNrf2 proteome may change the redox status through the formation of mixed disulphide bonds, which will then lead to irreversible protein aggregation in the myocardium. Of note, multitude of steps between transcription and translation may provide different regulatory or pathological check points in these hearts, which needs further investigation.

Our results presented here may provide clues to investigate whether the PTMs in each protein might have direct impact on enzyme kinetics and indirect influence on metabolomics of RS cardiomyopathy. However, the intensity or the frequency of altered enzyme kinetics due to the proteomic modifications is not explored in this study. For the first time, we report that a reductive myocardium displays over 50% mismatch between the mRNA and protein levels, warranting a high-throughput transcriptome, proteomic and molecular approaches to mechanistically understand the impact of RS. Our results will facilitate directions for future investigations on molecular signaling of RS and to identify novel bio-diagnostic-markers for reductive stress.

### Translational impact

Activation dependent dual roles of Nrf2 in maintaining redox balance is very crucial for the myocardial redox proteins. We hypothesize that over-the counter antioxidants intake recapitulates the caNrf2 mouse model used for the proteome analysis. Long-term consumption of these antioxidants will lead to progressive change in redox arm towards reductive side, thus a reductive stress myocardium. In a RS milieu, we observed downregulation of several myocardial adaptational/rescue pathways and upregulation of pathophysiological pathways which leads to reductive stress cardiomyopathy over time (**Fig 7**). Nevertheless, the cumulative data provide a compelling rationale to develop personalized antioxidant therapeutic strategies to correct the redox abnormalities.

### Limitations of the study

We observed modifications for amino acids in N and C terminal in the entire peptide fragment in the core proteome, that might get modified during the processing/labelling of the protein with the TMT reagents. Even though we observed peptide fragment level modifications like oxidation, methylation, acetylation and NEM modification, TMT analysis is insufficient to precisely measure the level of distribution and the intensity of the modifications.

## Ethics Statement

This study was carried out in accordance with the recommendations of the Guide for the Care and Use of Laboratory Animals of the National Institutes of Health. The protocol was approved by the Institutional Animal Care and Use Committee (IACUC) at the University of Alabama at Birmingham.

## Author Contributions

The study was designed by NSR. TMT analysis were performed by CLD and KP. NSR and SS interpreted the data and wrote the manuscript. All authors read and approved the final version of this manuscript.

## Consent for publication

All authors verified the content and approved the final version for submission and publication

